# Fast Ion Beam Inactivation of Viruses, Where Radiation Track Structure Meets RNA Structural Biology

**DOI:** 10.1101/2020.08.24.265553

**Authors:** B. Villagomez-Bernabe, S. W. Chan, J. A. Coulter, A. M. Roseman, F. J. Currell

**Affiliations:** The Dalton Cumbrian Facility and the Department of Chemistry, The University of Manchester, Westlakes Science & Technology Park, Moor Row, Cumbria, CA24 3HA, UK; School of Biological Sciences, Faculty of Biology, Medicine & Health, The University of Manchester, Manchester Academic Health Science Centre, The University of Manchester, Michael Smith Building, Oxford Road, Manchester M13 9PL, UK; School of Pharmacy, Queen’s University Belfast, 97 Lisburn Road, Belfast, BT9 7BL

## Abstract

Here we show an interplay between the structures present in ionization tracks and nucleocapsid RNA structural biology, using fast ion beam inactivation of the severe acute respiratory syndrome coronavirus (SARS-CoV) virion as an example. This interplay is one of the key factors in predicting dose-inactivation curves for high energy ion beam inactivation of virions. We also investigate the adaptation of well-established cross-section data derived from radiation interactions with water to the interactions involving the components of a virion, going beyond the density-scaling approximation developed previously. We conclude that solving one of the grand challenges of structural biology — the determination of RNA tertiary/quaternary structure — is intimately linked to predicting ion-beam inactivation of viruses and that the two problems can be mutually informative. Indeed, our simulations show that fast ion beams have a key role to play in elucidating RNA tertiary/quaternary structure.

## Introduction

At the time of writing the world is gripped with its fifth global pandemic since the Spanish flu outbreak of 1918 [1]. During the last decade viral epidemics have become a prevalent global health threat requiring techniques for rapid vaccine development [2]. Radiation has long been used to inactivate virions [3] although the uncorrelated nature of the ionization produced by gamma rays means that the protective outer envelope of the virus is highly damaged. High energy ion beams however produce long-thin (nanoscale) damage tracks which can be used for much more selective virion inactivation [4]. Currently, in an important extension of this concept, a large-scale ion beam facility, is using beams of heavy, high energy ions to attempt to produce a vaccine for SARS-CoV-2 [5]. Since the ions form long straight damage tracks, each of the ions interacting with a single virion will produce many ionizations or excitations in the nucleocapsid. Each of these events has a chance to damage the RNA, rendering the virion inactive. Damage to the envelope is typically limited to one or two locations, on the ion’s ‘way in’ and on its ‘way out’, thus limiting the amount of structural damage to the surface spike proteins integral to activation of a host immune response. RNA is a highly serial structure in its biological information expression; damage it in one or a few critical places and you probably render the virion inactive. In contrast the envelope proteins and phospholipid membrane are highly parallel structures so several of them can be radio-damaged without affecting the structure of the envelope significantly, leaving its function unchanged. This assertion points to an informatic aspect of the problem, one we will investigate in this paper - there are two structures involved, the structure of the virion being irradiated and the structure of the radiation tracks doing the irradiation. As we show below, the interplay between these two structural forms is a governing factors determining the predicted dose inactivation curve for any given virion.

If one can thus inactivate sufficient numbers of virions, they could be used as the key component for developing viral vaccines in a universal manner. The inactivation process, perhaps uniquely, produces little or no apparent change to the structure of the virions’ envelopes. Hence, they can be expected to enter a host in the usual way, including, in the case of SARS-CoV-2, using angiotensin-converting enzyme 2 (ACE2)-mediated cellular entry [6]. Upon encountering host immune cells, these inactivated antigenic virions would stimulate an immune response in the usual manner, but critically would lack reproductive capability. Hence, we call virions modified in this way Structurally Intact Radiation Inactivated Virions (SIRI-Vs).

It is worth asking the question “What would the specifications of an ion beam facility designed to produce sufficient quantities of SIRI-Vs look like?”. High-energy, large scale ion facilities such as the one currently being used [5] are rare and very expensive. However, we believe it is also possible to use mid-range ion facilities, such as those more traditionally associated with nuclear energy research [7].

In order to support such endeavors, it is important to have a set of predictive tools able to determine dose-inactivation curves reliably. Here we show rapid prediction of inactivation responses and discuss some of the barriers which must be overcome to produce reliable predictions. We believe it is currently premature to rely on radiation transport simulations such as those in [5] or those presented here. This does not mean they are unimportant. Instead a concerted, community effort is required to develop tools able to support calculations of this type. It is in this spirit that several new approaches are offered in this paper.

Even before the current coronavirus disease 2019 (COVID-19) pandemic, total human vaccine business was projected to be worth ~$60 Bn annually by 2024 [8] whilst that for animals is projected to reach ~$11Bn by 2025 [9]. Clearly then there would be commercial benefit in developing mid-range ion facilities similar to [7] but designed specifically for purpose.

## Results

### Fast ion irradiation of the SARS-CoV virion

A putative model of the SARS-CoV virion was developed using available structural biology information [10–13]. We chose to do this for SARS-CoV rather than the more topical SARS-CoV-2 since the structural biology database is more mature, allowing us to work with a realistic putative geometry. There is important information about the tertiary structure of at least part of the RNA in the nucleocapsid [10, 11]. The two virions are very similar so general conclusions reached should also apply to SARS-CoV-2.

The full details of the RNA tertiary/quaternary structure within the nucleocapsid are unknown. To test the importance of this unknown structure we considered three extreme forms of RNA arrangement, without making reference to any published structural forms. These structural forms were i) all RNA concentrated in a small sphere at the centre of the nucleocapsid, ii) all RNA concentrated in a spherical shell at the inner surface of the nucleocapsid shell and iii) all RNA uniformly and randomly distributed throughout the entire volume of the nucleocapsid and with no persistence from ion to ion. In each case the same total amount of RNA was included, as dictated by its sequence [14]. Whilst forms i) and ii) are obviously extremes, it is not immediately obvious that form iii) represents an extreme. Indeed, this form has been implicitly assumed in modelling inactivation of SARS-CoV-2 [5]. However, this form represents an extreme when viewed informatically as it places the tertiary/quaternary structure of RNA in its highest entropic state, something not typically found in biology. The whole field of structural biology derives much of its relevance from the fact that biological systems assume highly ordered structures, sometimes at considerable energy cost, in order to dictate function. This general tenet is ignored in assuming the RNA takes up such a high-entropy conformation. Furthermore, it is known that much of the RNA-protein complex of the SARS-CoV virion is highly ordered [10, 11]. Accordingly, SARS-CoV-2 cannot have its RNA in such a high entropic state.

For the geometry shown in figure 1, many 3 MeV He^2+^ ions were successively passed through the virion each entering the simulation at a different random location. The location of ionisation events were recorded, with ~42% of events being direct He^2+^ ion-interactions with the remaining ~58% being due to child electrons produced upstream. The range of initial positions of the ion trajectories was chosen to be significantly bigger than the area of the virion presented to the ion beam so that roughly 50% of the ions had no interaction with the virion, properly mimicking the effect of ions passing in close proximity to the virion causing damage to the membrane or spike proteins. This damage is generated through the creation of electrons which travel several nm from the main ion trajectory before causing an ionisation event. An example of this phenomenon is track 2, figure 1. An ion was considered to have interacted with the virion if even a single ionisation process occurred anywhere in the virion geometry. Hence, proper weighting was given to the effect of peripheral ion trajectories on the outer parts of the virion structure. Only ionisation events were used for this process, rather than energy deposition events as was done previously [5]. Although energy deposition approaches have been very successful in predicting biological outcomes in other nanoscale radiobiology contexts [15–17], we argue in the discussion section that more of a target-theory inspired approach [3, 18] is appropriate, although we admit this is an open question and one worthy of further investigation. However, none of the conclusions reached regarding the role of RNA tertiary/quaternary structure depend on the choice between energy deposition or target theory approaches.

**Figure 1.**
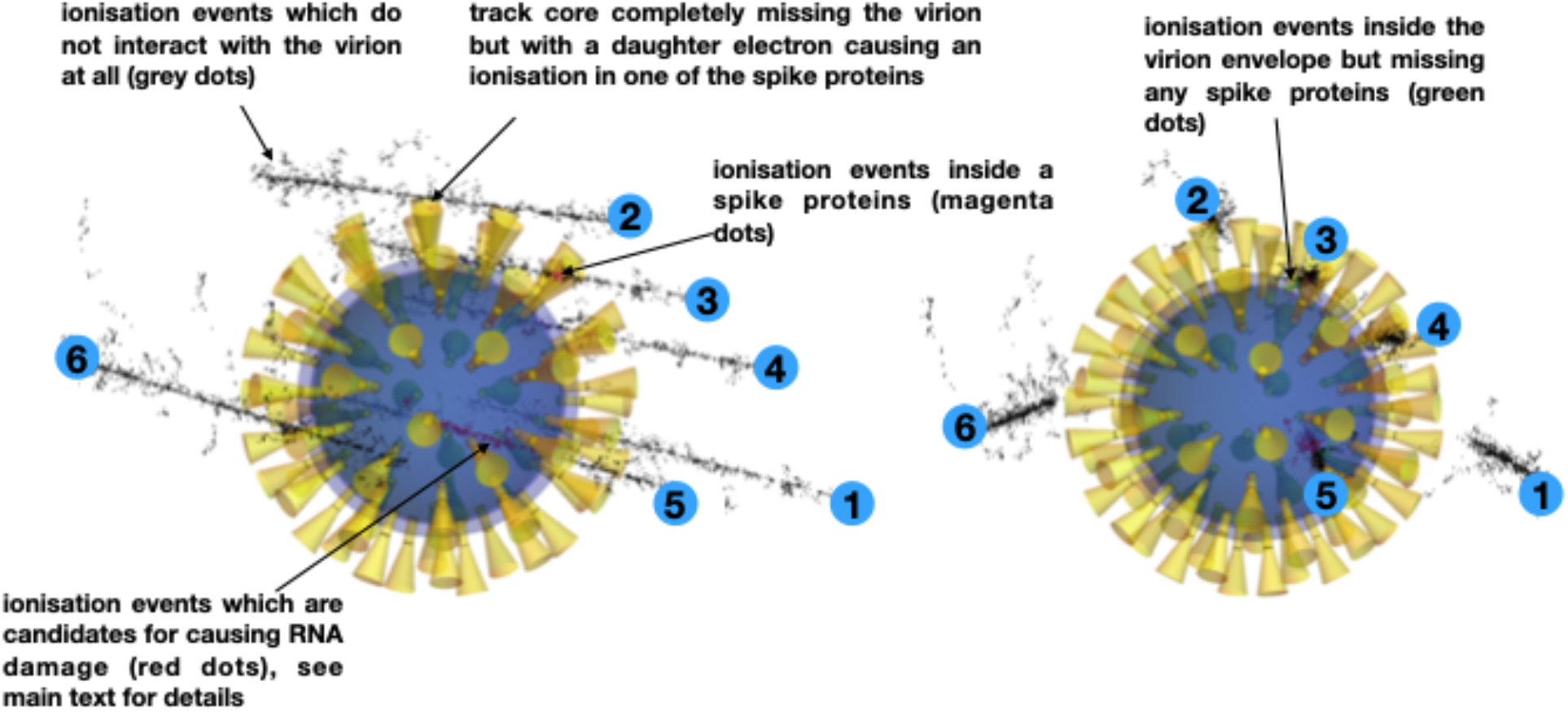
Geometry of the SARS-CoV virion and its interaction with six randomly generated 3 MeV He^2+^ ions shown from two different viewpoints. Tracks 1 and 6 completely miss the virion, whilst the core of track 2 misses but one of the child electrons produces an ionisation event in one of the spike proteins. The cores of tracks 3 and 4 pass through a spike protein, whilst 5 passes through the nucleocapsid causing RNA-candidate damage events (red dots), envelope damage events (green dots) and spike-damage events (magenta dots). It is necessary to include tracks such as 1 and 6 with cores far from the nucleocapsid to ensure a correct weighting of the spike-damage, i.e through tracks like 2. Full details of the geometry and how it was derived are given in the Methods section.

There are clear differences in the probability of inflicting damage to the RNA. In case i), the RNA presents the smallest area to the incoming ion beam, so it is not surprising to find the probability of no ionisations in the RNA to be significantly larger than in the other two scenarios. Although scenarios ii) and iii) have apparently similar histograms on a linear scale, there are noticeable differences. When examined on the logarithmic scale the histogram of the number of hits to the RNA shows marked differences in all 3 cases. Assuming a single ionizing event to the RNA is sufficient to inactivate a virion, then the key parameter is the probability of no ionizing events occurring within the RNA per ion track interacting with the virion. This probability has a value of 0.88, 0.50 and 0.52 for cases i) – iii) respectively. From these probabilities one can deduce a dose-inactivation curve using Poisson statistics [5]. However, it is not yet clear how much of the RNA constitutes the critical target, another factor which will also affect this curve.

Figure 3 shows histograms recording the average number of hits to spikes per ion track and the total number of spikes hit per ion track. These histograms are the same regardless of the RNA tertiary/quaternary structural form so only one set are shown. Unsurprisingly, the forms of the histograms for number of hits per spike and number of hits to the RNA in case ii) are similar since they both involve the ion track interacting with a hollow, essentially spherical structure. Typically, either 1 or 2 spikes are hit per ion, as might be expected since an ion has a chance to interact with spikes on its way in and on its way out. This result confirms that shown in [5] and is encouraging for the field of ion-beam inactivation of virions.

### Radio-density scaling

In order to differentiate between RNA and proteins, one must account for the differences in their interactions when subject to fast charged particles. Here, results of an analytical analysis of the scaling properties are presented and used to generate either scaled effective volumes or scaled effective densities for the biomolecules concerned.

Because there are not yet a good set of well curated charged-particle interaction cross sections for either proteins or RNA, one must rely on a well curated set for water. One particularly useful and reliable set are those in Geant4 DNA. One can simulate other biological materials by scaling the density of Geant4 water [19, 20] to that of the material concerned, as was done in [5] and in producing the data shown in figures 2 and 3. However, it is not *a-priory* clear that the mass-density is the appropriate parameter when scaling from water. The fast ions are really interacting with charge, not mass, as is evidenced by the Bethe formula [21, 22], which for non-relativistic ion velocities reduces to

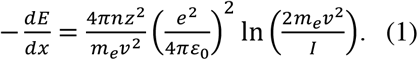

**Figure 2.**
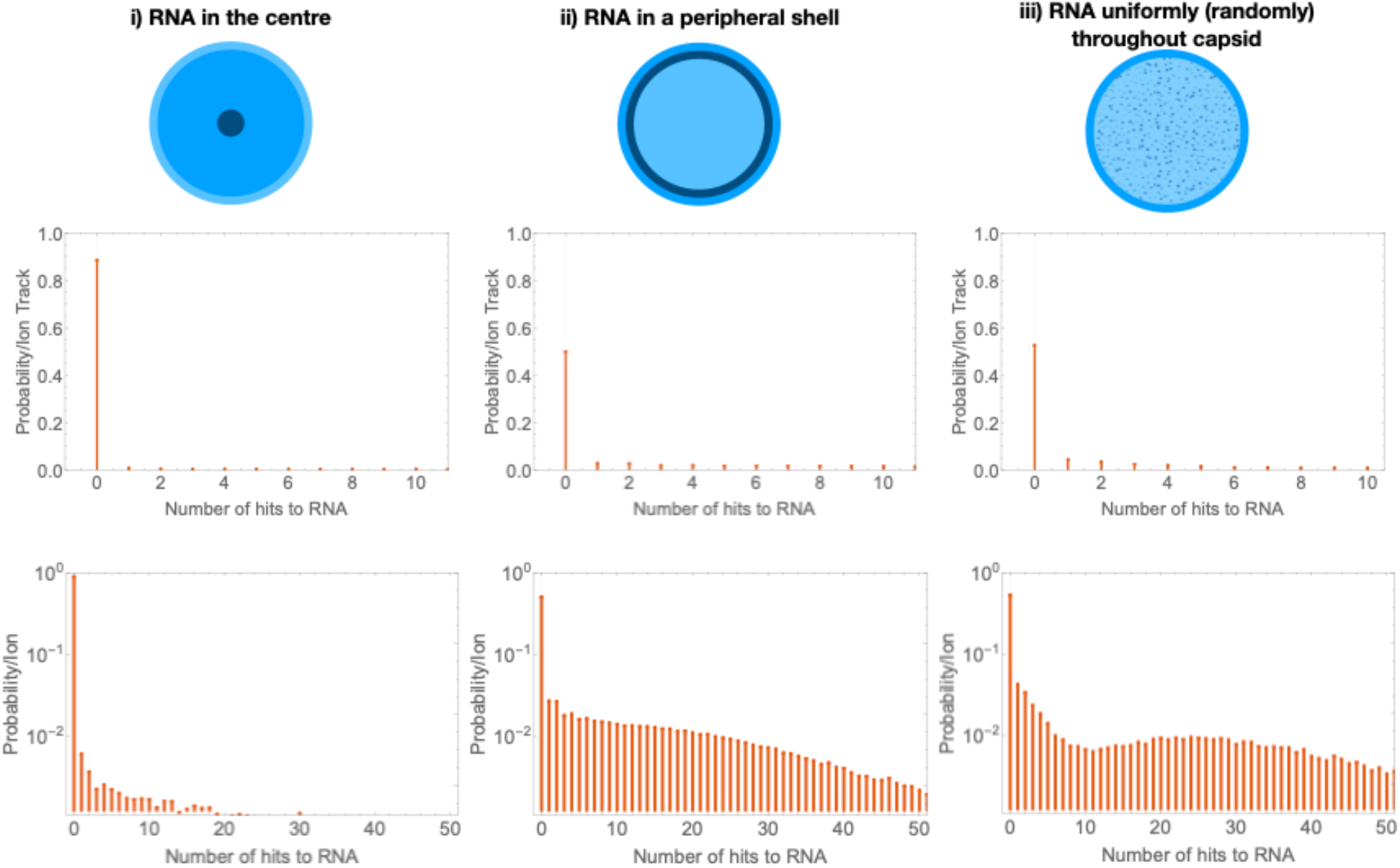
RNA distribution affects propensity for ionisation interactions. The top row symbolically shows the RNA configurations under consideration (column by column). The second row shows histograms of the probability of inflicting a number of ionising events to the virion’s RNA per ion track interacting with the virion on a linear scale whist the bottom row uses a logarithmic scale to show the long tail involving multiple hits to the RNA.

**Figure 3.**
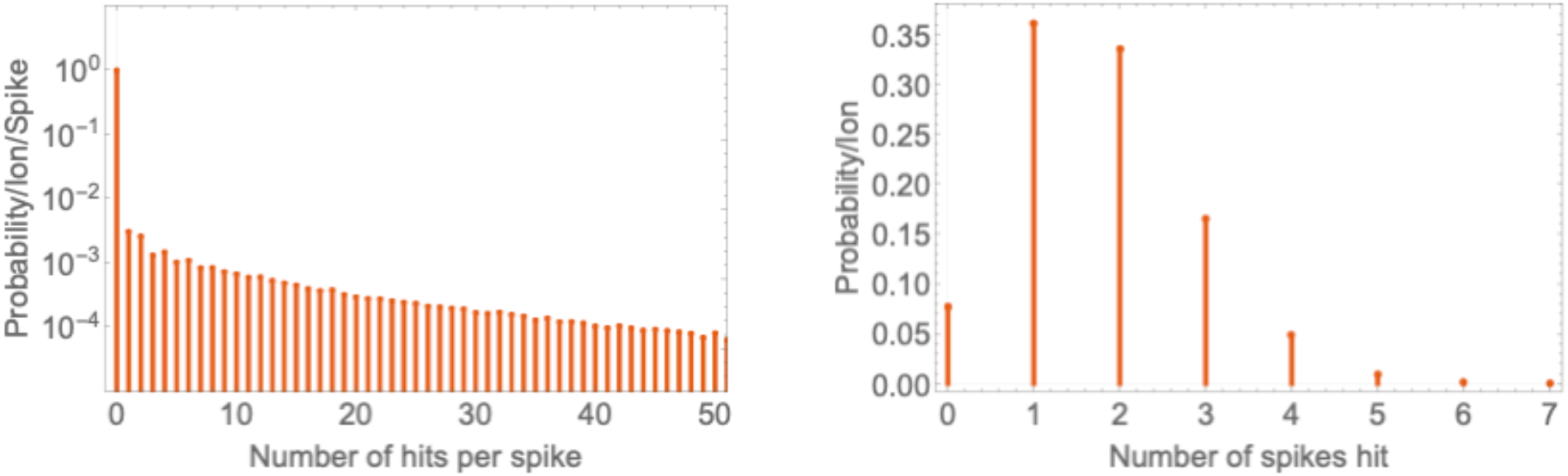
Probability of ionisation interactions with spike proteins. Histograms of (left) the probability of any selected spike protein complex being subject to an ionising event per ion track interacting with the virion and (right) the number of spikes undergoing an ionizing event per ion track interacting with the virion.

Here *ν* is the velocity of the ion, *z* is its change in multiples of the electron charge, *n* is the electron density of the material through which the ion is passing and *I* is the mean excitation potential. All other symbols take their usual meanings. The electron density is given by

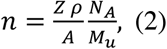

where the material dependent parts are *Z*, its atomic number, *ρ*, its mass density and *A*, is its relative atoms mass. The other two symbols are *N*_*A*_, the Avogadro number and *M*_*u*_, the molar mass constant. One can calculate the electron density for a compound as if it was simply a mixture of the atoms, i.e. one sums over all of the atoms present, replacing the density by the density of that atom in the material.

For most light atoms the ratio *Z*/*A* ≈ 1/2, allowing factorization of *Z*/*A* out of the expression for the compound electron density, providing a scaling directly between charge and mass density. However, hydrogen is a notable exception where the ratio *Z*/*A* = 1. Hence more hydrogen-rich materials will have proportionally greater electron densities and hence will interact with charged particle beams more intensely.

The other material-dependent term in equation (1) is 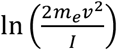. It has been shown that a range of biomolecules including water, proteins and RNA all have a mean excitation energy of approximately 70 eV [23]. Hence, this term can be ignored in scaling from charged interactions of water to those of other biomolecules. This argument can also be applied to the child electrons; hence one can find a set of scaling densities which can be applied to water, so it has approximately the same interaction rate as the material of interest. Taking known elemental compositions of biomolecules, it is then possible to generate a set of radio-equivalent densities such as those of table 1. These radio-equivalent densities are the density water would have to have to have the same electron density and hence the same rate of energy loss as per equation 1.

**Table 1.** showing the elemental compositions of various biomolecules, their densities and the resultant scaled radio-densities, i.e. the density water would have to have to produce the equivalent material-dependent term in equation 1. Elemental compositions are taken from [24].

|  | fractional density |  |  |  |  |  |  |  |  |
| --- | --- | --- | --- | --- | --- | --- | --- | --- | --- |
| Bio-material | H | C | N | O | P | S | density [g/cc] | relative electron density | Scaled radio-density [g/cc] |
| water | 0.111 | 0 | 0 | 0.889 | 0 | 0 | 1 | 0.555 | 1 |
| Protein C | 0.07 | 0.54 | 0.17 | 0.2 | 0 | 0.027 | 1.41 | 0.758 | 1.367 |
| Rubisco | 0.07 | 0.54 | 0.17 | 0.21 | 0 | 0.012 | 1.41 | 0.755 | 1.360 |
| Myosin | 0.07 | 0.53 | 0.17 | 0.22 | 0 | 0.008 | 1.41 | 0.752 | 1.355 |
| Collagen | 0.07 | 0.54 | 0.18 | 0.21 | 0 | 0.005 | 1.41 | 0.757 | 1.364 |
| RNA | 0.03 | 0.36 | 0.16 | 0.35 | 0.1 | 0 | 1.75 | 0.898 | 1.618 |

If the density of the biomolecule is not known in the particular configuration of interest, usually both its elemental composition and total mass are known, e.g. from primary sequence information. The density of the biomolecule can then be estimated from atomic volumes to generate a mass-density. Hence results like those of table 1 can be generated for many biomolecules as required.

The choice to scale the densities is somewhat *ad hoc*, indeed there is an assumption being made about the volumes of the objects which is used to determine their mass-densities. Since these are small objects compared with the length-scales over which the ion slows down one could fix the densities of the objects representing the various biomolecules in a simulation and instead scale their volumes, i.e. instead of slightly perturbing the densities of objects in a simulation to account for their different interactions with charged particles, one could instead slightly perturb their volumes. This second approach has the advantage that the entire nanoscale radiation transport simulation can now be performed at a constant mass-density. A set of ion tracks can be calculated for one density and then be ‘replayed’ many times with different starting positions and angles. Because the tracks do not pass through regions of different densities, the computationally expensive boundary-crossing checks are not required, instead one simply has to consider the volumes in which the transformed ionisations occur to determine the effects of the track.

We have explored both approaches (i.e. density and volume scaling) in the next section where we present results for ions passing through the RNA-protein nucleocomplex of the SARS-CoV virion. The scaled radio-density approach and the scaling of volumes instead of densities appear to be general procedures which can be applied in the calculation of radiation transport through nanoscale biological systems and they will be subject of a future publication.

### Simulations of the RNA-protein complex

In order to begin to assess the effect of RNA tertiary structure, we created a geometrical model of the RNA nucleocapsid complex for which there exists a putative RNA structure. The geometry of the simulation was drawn from the putative structure for the RNA-protein complex [10, 11]. Of course, the structures presented in these papers are far more sophisticated than the geometry used in our simulations – our geometry was simply constructed to capture the salient radiation biology effects. The RNA forms two helices separated and held in place by a protein complex, with the sugar-phosphate backbone lying on the inside of the helix and the RNA bases facing outwards. These two separate RNA strands are about 5 nm apart and their bases are not paired. The volumes and densities of the RNA bases were chosen in 3 ways. In the first set of simulations the entire volume was treated as having a density of 1.40 g/cm^3^, close to the asymptotic value for large proteins [25], and the volumes of the RNA bases and phosphate backbone were taken from [26]. This is equivalent to transferring the assumption made in [5] and our simulations above that the density is the same across the entire capsid down to the scale of the RNA-protein complex – clearly this is wrong but it serves as a useful starting point.

Simulations were performed in two ways, one where a full Geant4 DNA calculation was performed, and the locations of the ionisations was recorded. In a second simulation a group of tracks were replayed many times having been translated to enter the simulations at random places. The locations of the ionisations were scored using Mathematica’s geometry primitives. In both cases the density was set to 1.40 g/cm^3^ throughout.

In a second pair of simulations the density for the RNA was calculated using appropriate atomic volumes [26] so that the RNA backbone had a density of 1.83 g/cm^3^ and the bases had a density of 1.65 g/cm^3^. This simulation did not take account of the specific bases in the sequence as the motif being simulated is a repeating unit throughout the capsid, accounting for containment of more than half of the RNA complex [10, 13]. Instead the known RNA sequence [14] was used to calculate an average volume and density for the nucleobases. These densities were used in a second Geant4 DNA simulation otherwise similar to the one described above, whilst the same densities were used to scale volumes in a simulation which again replayed many tracks and used Mathematica’s geometry primitives to determine which ionisations occurred within the RNA backbone and base structures. Here the volumes of the bases and backbone elements were scaled by a factor of 1.65/1.40 and 1.83/1.40 respectively so that each element of the simulation had the same probability of interacting with the radiation.

In a third set of simulations, the same procedures were carried out using scaled radio-densities taken from table 1. Again, this was used to scale the density in a full Monte Carlo simulation and also to scale the volumes in a simulation where many tracks were replayed. The results of these simulations are presented in table 2. These simulations were conducted with the ions only coming in from one direction. In all cases, the fraction of ionisations found in simulation was greater than expected on purely geometrical grounds, suggesting the localised structure of the RNA does play a role, i.e. the probability is 5-10% greater than would be expected if transferring an approach like case iii) for the whole virion down to the scale of the RNA-protein complex. Since this result shows it is not appropriate to apply the approximation used in case iii) to the RNA-protein complex shown in figure 4 and given that approximately half of the RNA is in these RNA-protein complexes, it follow that this approximation made by us in case iii) and also used in [5] is not valid. Although they are generally in close agreement, the scaled volumes approach (scored using Mathematica) produces consistently smaller values than the scaled density approach calculated using Monte Carlo radiation transport. This could be due to differences in the volume-scoring algorithms when applied at sub-nm scale and will be the subject of further investigation.

**Table 2:** Volumes and densities used for the various simulations of the RNA-protein complex. F denotes the fraction of ionisations, with sim denoting the value obtained from the simulation and geo denoting the fraction expected on purely geometrical grounds, i.e. the fraction of the density-volume product occupied by either the base or backbone components. Typical statistical errors associated with the simulated Fs were between ±0.005 and ±0.01 × 10^−3^.

| Method | V<sub>base</sub><br>[nm<sup>3</sup>] | ρ<sub>base</sub><br>[g/cm<sup>3</sup>] | F<sub>base</sub><br>sim/10<sup>-3</sup> | F<sub>base</sub><br>geo/10<sup>-3</sup> | V<sub>bbone</sub><br>[nm<sup>3</sup>] | ρ<sub>bbone</sub><br>[g/cm<sup>3</sup>] | F<sub>bbone</sub><br>sim/10<sup>-3</sup> | F<sub>bbone</sub><br>geo/10<sup>-3</sup> |
| --- | --- | --- | --- | --- | --- | --- | --- | --- |
| Average Density M/C | 0.13 | 1.40 | 0.57 | 0.52 | 0.18 | 1.40 | 0.76 | 0.71 |
| Average density Mathematica | 0.13 | 1.40 | 0.55 | 0.52 | 0.18 | 1.40 | 0.78 | 0.71 |
| Mass-density-scaled densities M/C | 0.13 | 1.65 | 0.68 | 0.61 | 0.18 | 1.83 | 1.02 | 0.93 |
| Mass-density scaled volumes | 0.15 | 1.40 | 0.65 | 0.61 | 0.23 | 1.40 | 1.00 | 0.93 |
| Charge-density scaled densities M/C | 0.13 | 1.54 | 0.65 | 0.57 | 0.18 | 1.70 | 0.98 | 0.87 |
| Charge-density scaled volumes | 0.14 | 1.40 | 0.62 | 0.57 | 0.21 | 1.40 | 0.94 | 0.87 |

**Figure 4.**
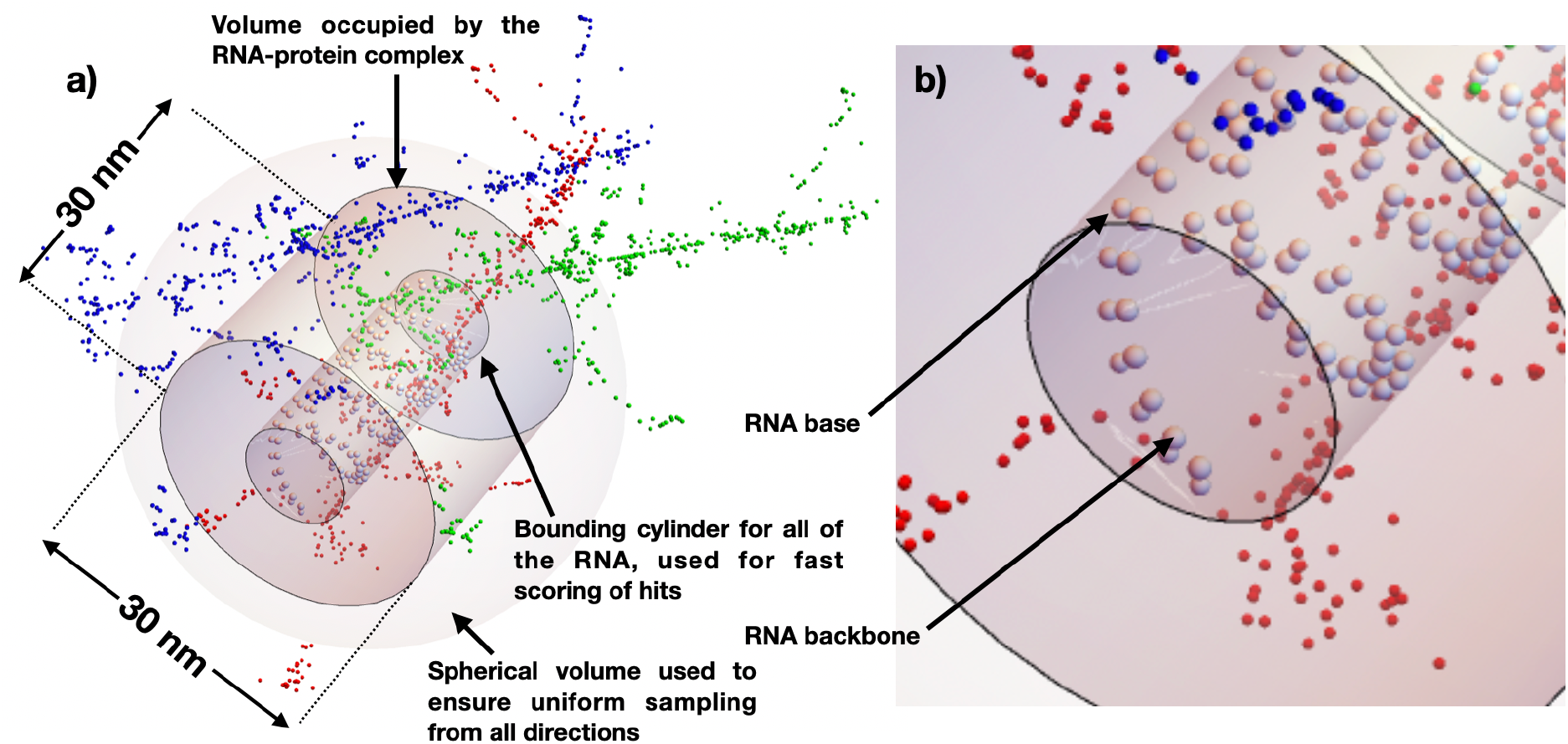
nucleocapsid protein-RNA geometry showing three separate tracks due to 3 MeV He^2+^ ions (red, green and blue), showing the various volumes used to define the simulation geometry.

Unlike the virion as a whole, the protein-RNA complex does not have quasi-spherical symmetry, so a full simulation should average irradiation over all orientations although this is computationally more expensive. Using volume-scaling with scaled radio-densities, we performed another simulation. Here we not only scored hits to the various RNA structures; we also stored the full track identification for each ionisation within any of the RNA structures. The simulation geometry is shown in figure 4.

These were then analysed for coincident events, i.e. when a single primary ion caused ionising events in two different parts of the RNA structure. The results are shown in figure 5. Unsurprisingly, most of the coincidental ionisation events concern proximal backbone/base units, i.e. where the difference in index (explained in figure 5’s caption) is < 5. However, it is interesting to note that there are persistent features in this map, even when structures are several nm apart. The most prominent features are found when the difference in indexes is 32 or 53, which corresponds to the kinds of coincident events shown by the pink arrows in figure 5a). However, there is not a feature found corresponding to a difference in index of 44, i.e. a pair of events on directly opposite units of the two strands. There is also a weak shoulder on the same-strand distribution corresponding to a difference in index of about 17 although there is no immediately obvious cause of this feature. If each ionizing event corresponds to a chance to break the RNA strand where the ionisation took place, it follows that, depending on details of how the RNA is ordered within the nucleocapsid, the lengths of strands produced by multiple events from a single ion will reflect features of the map shown in figure 5b), i.e. they will reflect the tertiary RNA structure. This idea is discussed in more detail in the Discussion section.

**Figure 5.**
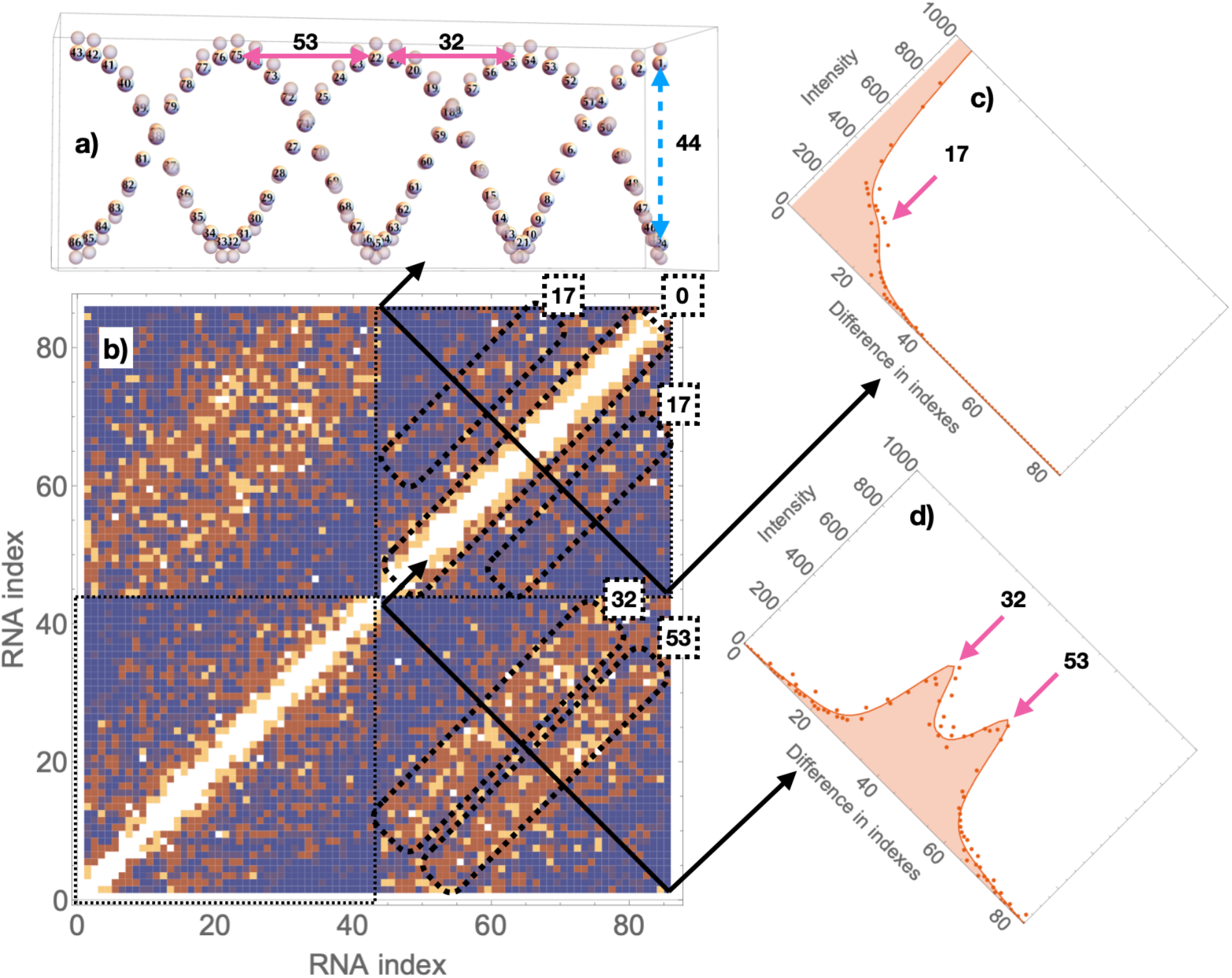
Plot showing the effect tertiary RNA structure has on coincidental ionisation of two separate parts of the RNA structure by a single ion. Panel a) shows the indexing system used to denote locations within the two helices of the RNA. The index increases from 1 to 43 running along one strand and then (right to left as shown here) and then back to the rightmost nucleotide of the opposite strand running 44 to 86. Pertinent differences in indexes are illustrated by the labelled double-headed arrows. Panel b) shows a 2-D map of the number of times two separate ionizing events happen on the backbone/base combination from the same ion as a function of the indexes at which the two events occur. Note this plot contains 4 quadrants, those bottom-left and top-right corresponding to two events on the same helix, those top-left and bottom-right corresponding to two events, one on each helix. Panels c) and d) show the projections of these plots, summing all features with a constant difference together. The dots represent the sums over the 2-D map (i.e. the Monte Carlo data), the solid lines represent predictions made using our ‘Method of Ionizing Lines’ described in the main text.

### Method of ionising lines (MoILs)

Another fast analytical technique was developed, this time to account for the spectrum of coincident events from single ion tracks causing ionising events in two separate parts of the RNA structure, with a view to developing a framework for rapidly assessing the likely effect of tertiary structure on two-site damage to various RNA structures. Every pair of structures was considered in turn with a histogram constructed with the bins being the difference in position along the strand of the pair and the contribution to the height being 1/*r*^2^, where *r* is the Euclidean distance between the two structural elements. This factor acknowledges the fact that an ion track is long and narrow so that the chance of it interacting with two structures of fixed volume is proportional the solid angle one of the structures presents to the other and hence proportional to the inverse square of their separation. This method could be a cornerstone of an approach to critique RNA structures in general. As can be seen in figure 5c) and d) this method is able to rapidly predict the distribution of coincident ionizing events as a function of index difference, i.e. for a suitable RNA ordering within the nucleocapsid it is able to rapidly predict the strand-length distribution for coincident single-strand breaks. This concept is considered further in the discussion section.

## Discussion

The histograms presented in figure 2 clearly show that 3D arrangement of the RNA (i.e. the tertiary/quaternary structure) within the virion is important. Different assumed tertiary structures give different probabilities of delivering one or more ionisations to the RNA and hence give different probabilities of rendering a virion inactive per ion. In coronavirus most of the viral RNA lies within ~25 nm of the inner face of the membrane [12], making the arrangement similar to case ii). However, in this region it is believed to be largely in highly ordered RNA-protein complexes such as those shown in figure 4 [10]. As table 2 and figure 5 show, the probability of hitting the RNA in such a configuration is roughly proportional to the fractional volume occupied by the RNA, although it is consistently higher by 5-10% – something possibly related to the RNA’s tertiary structure. The separate RNA-protein complexes will keep RNA far apart meaning that the coincident damage probability across two of these complexes will be small. Taken together these factors suggest something close to case iii) would apply. However, the predicted inactivation probability would be raised somewhat due of ionising events on the RNA due to the structure in the RNA-protein complex. Hence, one might suggest that the actual dose inactivation curve might lie somewhere between that for cases ii) and iii) considered.

To make more firm predictions, among other things, one would need a better knowledge of the RNA tertiary/quaternary structure, something which is considered a major challenge of structural biology. However, it is not possible to deduce the tertiary/quaternary RNA structure from the dose-inactivation curve – there could be many different tertiary/quaternary structures which lead to the same dose-inactivation curve. Furthermore, there is currently significant uncertainty in radiation transport cross-sections, the probability of an RNA strand-break either per ionizing event (target theory) or the amount of energy required to make a break (energy deposition theory). These factors will need to be systematically addressed before meaningful dose-inactivation curves can be deduced, even for an assumed tertiary/quaternary RNA structure.

Historically there have been broadly two approaches to relate physical radiation transport processes to biological outcome, target theory [3, 18] and energy deposition [15–17]. Although it is an open question and one requiring further research, we argue that a target theory-based approach is more appropriate. The energy deposition approach accounts for energy deposition by excitation (of water) as well as its ionisation. In liquid water both processes lead to the formation of reactive products which mediate biological damage. However, in biological structures such as proteins and RNA, there is a larger network of bonds through which electronic energy can be transferred to stabilise the system, hence reducing bond breaking. In contrast, ionisation will almost inevitably lead to rapid bond breaking. Indeed, the system is more complex and might need an entirely new theory altogether. For example, it is reasonable to consider a highly folded 3D biological structure to be more robust to effects of ionizing radiation than a more open one.

We have presented some new tools for the rapid evaluation of the effects of ion beams on biological systems. The concept of volume-scaling rather than density-scaling needs further investigation although ultimately what is required is a set of cross-sections for atomic and molecular processes for RNA, DNA and proteins. The entire gamut of these biomolecules is clearly too large to study as a whole, representing the entire ‘Holy Trinity’ of structural biology. However, the charge-density scaling approach leading to table 1 is general, only requiring primary sequence information and atomic volumes. The approach, then, could form the basis of a new way of generating the required cross-section data as required.

In figure 5 we showed that there are preferred differences along the sequence for locations of coincident strand-breaks produced by a single ion. This was manifest as the differences in indexes which produced peaks in figure 5c) and d). Coincident ionisation events could lead to the RNA being broken in two places. Since the threading of the RNA through the SARS-CoV collection of RNA-protein complexes is not yet understood, it is impossible to say if these peaks in the differences of indexes will manifest as peaks in the differences of RNA strand-lengths observable after low dose irradiation (ensuring one or less ions ionise a given RNA strand) and disassembling of the capsid. If there were peaks in the strand-length distribution, this would then directly be informative of the RNA tertiary/quaternary structure in an entirely new fashion and in turn informative of the way the virion forms. This idea then points to the idea of using coincident RNA strand-breaks induced under single-ion hit conditions as a new means of deducing tertiary/quaternary RNA structure.

The method of ionizing lines (MoILs) presented here is completely general and allows for the rapid prediction of RNA strand-length distributions after irradiation in this manner. To illustrate this, we show an example of a prediction taken from the RNA-Puzzles collection, as is shown in figure 6. This clearly shows that the ion beam induced coincident fragment pattern is able to critique the best computationally derived tertiary structure from within the RNA-Puzzles collaboration compared to the ‘golden standard’ crystallography structure. Hence ion-beam induced fragment length distributions can provide clear indications of RNA tertiary structure. Essentially, the low-entropy pattern of ionisations induced by the ion beam is able to encode the tertiary structural information in the RNA in the fragmentation pattern.

**Figure 6.**
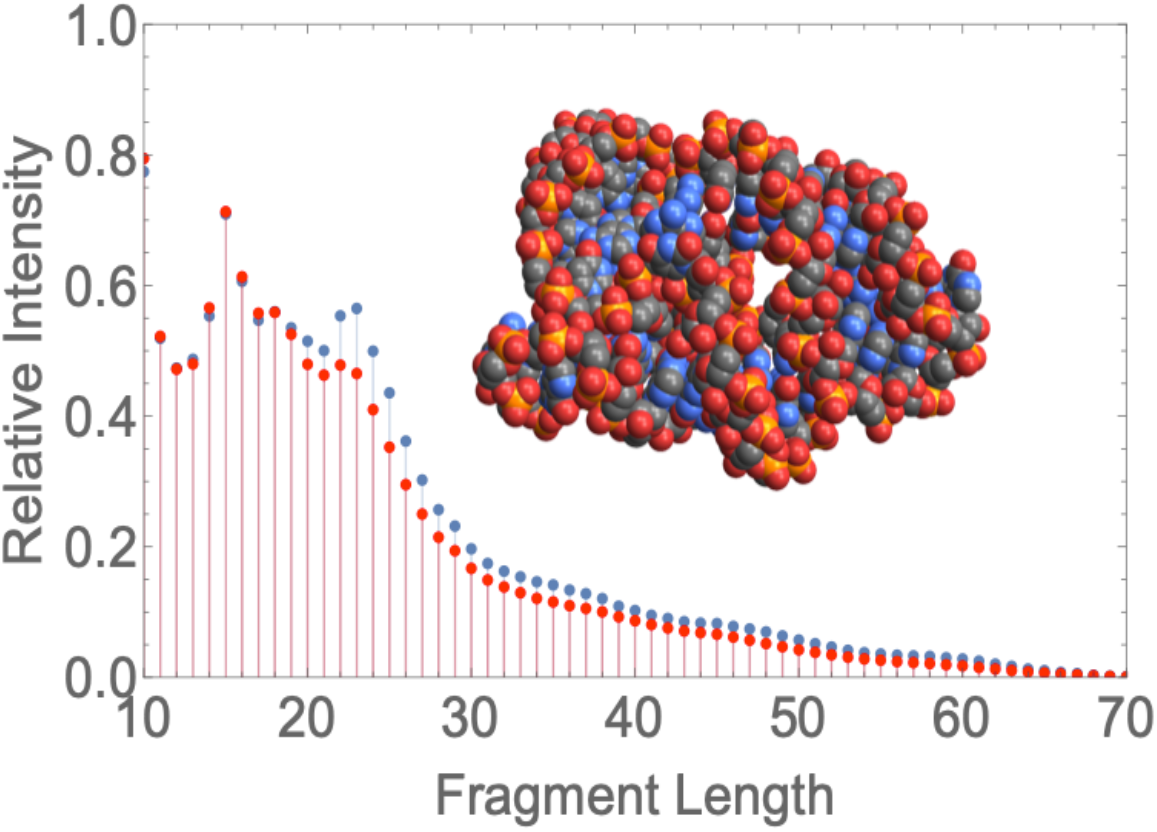
comparison of MoILs-predicted fragment length distributions of the best predicted (but not entirely correct) structure (blue dots on the histogram) and the ‘gold standard’ X-ray crystallographic structure (red dots on the histogram) for RNA puzzle #9 (a 71 nt RNA aptamer) from the RNA puzzles collection [27]. The X-ray crystallographic structure was also used to make the space-filling model shown in the inset. Note the peak at a fragment length of ~23 is indicative of incorrect structure prediction.

Features like this in the fragment length distribution could form the basis for critiquing and predicting RNA tertiary structures.

## Conclusion

We have shown that the interplay between radiation track structure and RNA’s 3D arrangement in space (i.e. the tertiary/quaternary structure) has a significant effect on predicted dose-inactivation curves for virions and that this effect can be traced back to the localised, low-entropy/highly informatic fashion ion beams transfer energy to biological systems. Indeed, it is this localised transfer which offers the promise of structurally intact radition inactivated virions (SIRI-Vs). This informatic transfer can potentially encode tertiary/quaternary structure structural information in the lengths of ion-beam induced RNA fragments, pointing to a new field of radio-structural biology. Methods such as charge-density based volume scaling of targets and MoILs have been proposed as tools to help develop this field with its promise of ab-initio RNA tertiary structure determination, provision of constraints for hybrid methods, detection of quaternary interactions and signatures to identify or monitor structures in-situ.

## Methods

TOPAS [28] version 3.3p01 that wraps the geant4-10-05-patch-01version [29] was used to transport 3 MeV alpha particles through a volume of water with a density of 1.4 g/cc over a distance of 0.8 μm. The slowing down of the ion is negligible over this distance. The vertices for ionization by either the primary ion or any child electrons were recorded on an ion-by-ion basis for future simulations. 100 such ion tracks were recorded and saved for future use. For the simulations shown in figures 2 and 3, one of this set of tracks was chosen at random and a 150 nm long section of the track was extracted. This extracted track was then shifted by a random amount in the x- and y- directions (the z-direction being the direction of propagation of the ion). The locations of the resulting ionization events were then recorded to produce the histograms shown in figures 2 and 3. For each of these, ionizations from 200, 000 independent ion tracks were recorded.

The geometry used for the Virion was derived from available structural biology information [10–13]. The inner and outer radii of the envelope were set to 40 and 44 nm respectively with the RNA being confined within a spherical region of radius 39 nm, subject to other conditions defining cases i) – iii) for the RNA secondary structure. A total of 75 protein spike complexes were distributed penetrating outwards through the virion’s envelope. Each spike complex was represented by an 8 nm long 1.6 nm radius cylinder starting from the inside of the envelope and pointing radially outward. The other end of the cylinder was joined to a truncated cone, exactly matching the 1.6 nm radius at the narrow end, widening 5 nm in radius at the wide end, also with its axis point away radially and also being 8 nm long. These spike complexes were positioned so that the axis of each one pointed directly away from the centre of the virion so as to be approximately equally spaced on the envelope’s surface. Each of these objects was represented by a Mathematica geometrical region, using Mathematica 12.1 with the RegionMember function being used to determine if an ionisation vertex was within a region of interest.

For simulation of the RNA-protein complex, again structural information was taken from [10–13]. The outside of the complex was constructed as a 30 nm long, 15 nm radius cylinder. Spiralling around the axis of this cylinder were two strands of RNA, each nucleotide being represented by two spheres, one of radius 0.35 nm (backbone) and one of radius 0.31 nm (base). The backbone was constructed along a 4.35 nm radius helix of pitch 14 nm with two complete revolutions of the helix being included for each strand. The bases were placed so as to just touch the backbone, with the axis joining them pointing directly outwards. This geometry was used for all the Monte Carlo (Geant4 [29]) simulations reported in table 1, with the RNA chain modelled as water spheres with densities as per table 1 and immersed in a protein environment modelled as a water with density of 1.4 g/cc.

Geant4 DNA [19] was used to track alpha particles and all secondary particles produced down to 10 eV and all deexcitation processes were included, such as Fluorescence, Auger cascade, and PIXE. The beam consisted of a rectangular field of 30 by 30 nm^2^ of 3 MeV mono-energetic alphas launched along an axis, perpendicular to the RNA chain. A TOPAS extension was implemented in order to retrieve all the pertinent information each time an ionization occurred such as position, energy deposited in the material, particle type that indicated the interaction, material where ionization took place, track ID, and event ID of the particle. The last two parameters allow to associate the track of all secondary particles that were created in the same event.

For the volume-scaled simulations, the volumes of the spheres representing the backbone and bases were modified as per table 1 with the scoring being done by replaying ion tracks and using Mathematica’s RegionMember function as described above. 200,000 ion trajectories were considered for each simulation. For the simulation shown in figures 4 and 5, the same process was repeated but the ion trajectories were rotated as well as translated so the sphere shown in figure 4 was uniformly and isotropically irradiated. The inner cylinder shown in figure 4 was used to speed up the simulations by using nested region checking since only ionisation events inside this cylinder can also be inside one of the spheres representing the RNA strands. 3.5 million ion trajectories were used in this simulation.

## Data availability

Once the manuscript is accepted all of the data used to produce the figures will be placed in Manchester University’s Institutional Repository as per the university’s policies to be publicly available and searchable.

## Author contributions

BVB performed the Monte Carlo simulations used throughout. FJC developed the Mathematica simulations, the charge-density and volume-scaling concepts as well as the Method of Ionizing Lines. He produced the manuscript and prepared all the figures. JC, SWC and AR provided advice on the biological systems being modelled. All authors reviewed the manuscript.

